# Yeast biopanning for detecting antibody binding to site-specific phosphorylations in tau

**DOI:** 10.1101/2022.01.15.476481

**Authors:** Monika Arbaciauskaite, Azady Pirhanov, Yu Lei, Yong Ku Cho

**Affiliations:** Department of Chemical and Biomolecular Engineering, University of Connecticut, Storrs, CT, USA; Department of Biomedical Engineering, University of Connecticut, Storrs, CT, USA; Institute for Systems Genomics, University of Connecticut, Storrs, CT, USA

**Author notes:** To whom correspondence should be addressed: Yong Ku Cho: Department of Chemical and Biomolecular Engineering, Institute for Systems Genomics, University of Connecticut, Storrs CT 06269. Grant numbers: NSF 1706743 NIH 1R21NS111358-01A1 Alzheimer’s Association 2019-AARG-NFT-640971.

**Keywords:** Tau, phospho-tau, post-translational modification, biopanning, antibody binding, yeast surface display

## Abstract

The detection of phosphorylated tau (p-tau) levels in clinical samples is of extreme importance for the detection of Alzheimer’s Disease (AD) as well as other neurodegenerative diseases. Recent reports show that detecting low levels of p-tau in plasma can be used as a reliable biomarker for detecting AD prior to the onset of memory loss. The ability to detect such low levels of p-tau is dependent on antibodies specific to the post translationally modified protein. However, the need for reliable phospho-site specific antibodies persists due to a lack of approaches for identifying monoclonal antibodies and characterizing non-specific binding. Here, we report a novel approach using the principles of yeast biopanning to create a robust platform that uses synthetic peptides as target antigens. Using peptides as antigens enables screening antibodies against defined post-translational modification sites, particularly for targeting intrinsically disordered proteins such as the human tau protein. To readily assess yeast binding and distinguish non-specific binding, we developed bi-directional expression vectors that allow antibody fragment surface display and intracellular fluorescent protein expression. We show that our platform can specifically and robustly detect a specific site within the p-tau target peptide when compared against non-phosphorylated controls. By improving biopanning parameters, we enabled phospho-specific capture of yeast cells displaying single-chain variable region fragments (scFvs) against p-tau with a wide range of affinities (K_D_ = 0.2 to 60 nM). These results demonstrate that yeast biopanning can robustly capture yeast cells based on phospho-site specific antibody binding, opening doors for facile identification of high-quality monoclonal antibodies.

## Introduction

Reversible protein phosphorylation plays a central role in how cells adapt to the environment, differentiate, and coordinate. Protein phosphorylation enables regulations at timescales much faster than gene expression, by inducing conformational changes that alter enzymatic activity, allosteric regulation, or protein-protein interactions (Graves & Krebs, 1999; Johnson & Lewis, 2001; Pawson, 2004). Abnormal phosphorylation is associated with many human diseases and is a source of toxicity in pathogens (Cohen, 2001). Phosphorylation patterns of the microtubule-associated protein tau is a prominent example, for which association with neurodegenerative diseases has been extensively documented. In tau, increased phosphorylation at specific sites shows strong correlations with disease stage (Wesseling et al., 2020) and progression rate (Dujardin et al., 2020). Moreover, elevation in plasma concentration of tau phosphorylated at threonine 181 or 231 (T181 or T231) is a strong biomarker of AD (Arbaciauskaite et al., 2021; Ashton et al., 2021; Janelidze et al., 2020; Thijssen et al., 2020). Therefore, there is a great deal of interest in accurate detection of site-specific protein phosphorylation.

Antibodies are widely used for detecting protein phosphorylation, but validation of their binding specificity remains a major challenge. High specificity phospho-site antibodies tend to interact with one or more phosphate groups on the modified residue and additional nearby epitopes to achieve site-specificity (Arbaciauskaite et al., 2021). Unfortunately, many existing phospho-site antibodies suffer from poor specificity, often showing cross-binding to non-phosphorylated target sites or other phosphorylation sites (Ercan et al., 2017; D. Li & Cho, 2020). Moreover, most available preparations are rabbit polyclonal, primarily because rabbits are highly immunogenic to small molecules and haptens unlike rodents (Y. Li et al., 2000; Liu et al., 2016; Weber et al., 2017). According to a curated antibody database (Labome), 93 % (15,234 out of 16,371) of existing phospho-site antibodies are rabbit immunoglobulins, and 89 % are polyclonal. Immunization with synthetic peptides that contain site-specific phosphorylations lead to reagents with broad applicability, but lack of monoclonal antibodies critically limits validation and reproducible detection of protein phosphorylation.

To address this challenge, we sought to develop a method for screening and characterizing antibody clones based on specific binding to phosphorylated target protein sites. Considering the challenge in finding high specificity clones, we aimed to achieve quantitative discrimination based on binding specificity. Here we report the implementation of yeast display biopanning with whole-well cell counting to measure phospho-site specific binding to the clinically relevant tau phosphorylation site T231. Based on the previous finding that yeast cells displaying an anti-fluorescein scFv can adhere to a monolayer of cultured mammalian cells chemically modified with fluorescein (Wang & Shusta, 2005), we predicted that yeast cells displaying phospho-specific scFvs can be biopanned. We show that synthetic phospho-peptides can be immobilized on a layer of human embryonic kidney (HEK) 293 cells and allow yeast biopanning based on specific antigen-scFv interaction. A bi-directional expression plasmid was introduced to enable scFv surface display and intracellular expression of fluorescent proteins in yeast. This allowed rapid cell counting using automated microscopy and the addition of in-well control yeast for detecting non-specific binding. We report improved peptide immobilization and biopanning conditions that allow phospho-specific capture of yeast displaying scFvs with a moderate affinity (K_d_ of 60 nM), well within the affinity level of antibodies identified from naïve libraries through yeast biopanning (Wang et al., 2007). The results clearly show that yeast biopanning can identify phospho-site specific antibodies and demonstrate the potential for clonal selection of phospho-site binders.

## Methods

### Bi-directional Expression for Yeast Surface Display of Antibody Fragments and Intracellular Production of Fluorescent Proteins

For bi-directional expression in yeast, plasmid vector pBEVY-GT (Addgene, RRID:Addgene_51231) containing the GAL1-10 promoter with two distinct terminators was used. Yeast enhanced green fluorescent protein (hereafter referred to as GFP) (Huang & Shusta, 2005) was cloned into the vector pBEVY-GT between restriction sites BamHI and PstI. pT231 tau scFvs variants (pT231 scFv, pT231 scFv mutants 3.24, or Y25A (D. Li et al., 2018)) were cloned into the vector pBEVY-GT between restriction sites XmaI and EcoRI, resulting in pBEVY-pT231 scFv (or other mutants)-GFP. To generate yeast expressing a control scFv, the anti-fluorescein scFv 4420 (a gift from Dr. Eric Shusta) (Boder & Wittrup, 1997) was inserted to pBEVY-GT using restriction sites XmaI and NheI. The scFv constructs contain a FLAG tag (DYKDDDDK) to its N-terminus (FLAG tag-scFv-Aga2p) to detect full-length scFv expression. To distinguish the control yeast, Golden gate assembly was used to insert a yeast codon mCherry (version 4) (Qian et al., 2012) into the vector pBEVY-GT, resulting in pBEVY-4420-mCherry.

Plasmid pAP208 was cloned for surface display of scFvs fused to GFP (FLAG tag-scFv-Aga2p-GFP). The Aga2p and secretion signal sequence were digested with restriction enzymes NheI and BsaI and the GFP sequence was digested with restriction enzymes BsaI and XhoI. The sequences were then cloned into the pCT-4RE backbone between restriction sites NheI and XhoI. All restriction endonucleases were purchased from New England BioLabs.

Resulting plasmid constructs were transformed into *S. cerevisiae* strain EBY100 (ATCC MYA-4941) (Boder & Wittrup, 1997) using frozen-EZ yeast transformation II kit (Zymo Research, Cat. No. T2001) and grown on SD-CAA agar plates for 3 to 4 days. From the plates, single colonies were picked and grown in 3 mL of SD-CAA medium at 30 °C with shaking at 250 rpm overnight. The cell concentration was then determined by measuring the optical density at 600 nm (OD_600_) and 10^7^ yeast cells were resuspended in 3 mL of SG-CAA medium at 30 °C with shaking at 250 rpm for at least 20 hours to induce expression of proteins on the surface.

### HEK293FT Cell Culture and Seeding

HEK293FT cells (Invitrogen Cat. No. R70007, RRID:CVCL_6911) were grown in Dulbecco’s Modified Eagle’s Medium (ThermoFisher Cat. No. 12320-032) supplemented with 10 % fetal bovine serum (Cytiva, head inactivated). The cells were used for less than 15 passages in continuous culture from cells that were previously frozen at early passage. Wells in a 96-well plate (Thermo Scientific Cat. No. 165305) were coated with 50 µL of Matrigel (Corning Cat. No. 354234) and the plate was rocked to ensure even coating. The plate was then incubated at 37 °C for at least one hour before cells were added. HEK293FT cells were seeded in the 96-well plate the day before experiments to reach 100 % confluency. The next day, the cells were used in the yeast biopanning protocol as described below.

### Yeast Biopanning against Tau Peptide Ligands

To carry out yeast cell biopanning against peptide ligands, HEK293FT cells are first grown to 100 % confluency in wells within a 96-well plate as described above. These wells were then washed twice with 100 µL of ice-cold PBSCMA (137 mM NaCl, 2.7 mM KCl, 10 mM Na_2_HPO_4_, 1.8 mM KH_2_PO_4_, 1 mM CaCl_2_, 0.5 mM MgCl_2_, and 1 g/L BSA, pH 7.4). Cells were then biotinylated using NHS-PEG_4_-Biotin (125 µM unless otherwise specified, ThermoFisher Cat. No. A39259) diluted in PBSCMA to achieve a total volume of 50 µL per well for 30 minutes at room temperature. After this, wells were washed again twice with 100 µL of ice-cold PBSCMA. To quench the biotinylation reaction, wells were incubated with 50 µL of PBSCM with 0.1 M Glycine for 10 minutes at room temperature. Wells were washed once with 100 µL ice-cold PBSCMA after this.

Streptavidin was then added to the cells at 50 µL total volume per well diluted in PBSCMA for 30 minutes at room temperature. For initial experiments aiming to visualize the degree of HEK293FT cell biotinylation, streptavidin conjugated with Alexa Fluor 647 (1:200, 155 nM, Invitrogen Cat. No. S32357) was used as the streptavidin reagent. For all other experiments, unconjugated streptavidin (1 mg/mL diluted to 155 nM unless otherwise stated, Sigma Cat. No. 85878) was used. After incubation, wells were washed twice with 100 µL of ice-cold PBSCMA.

The wells were then incubated with biotinylated peptide antigens (0.1 µM unless otherwise stated) diluted in PBSCMA to achieve 50 µL total volume per well for 30 minutes at room temperature. The peptides used were previously described biotinylated peptides (D. Li et al., 2018) containing a sequence found in the tau protein with and without phosphorylated threonine site pT231. The biotinylated phoshpo-peptide (KKVAVVR(pT)PPK(pS)PSSAK-biotin) and the non-phospho-peptide (KKVAVVRTPPKSPSSAK-biotin) were synthesized by Peptide 2.0. The phospho-peptide contains two phosphorylation sites, but the pT231 scFv interact only with the pT231 site (Shih et al., 2012). To test a range of peptide concentrations, the peptide was prepared to the highest concentration necessary for that day of experiments and then continuously diluted 2-fold to cover the range of the titration curve. Wells were then washed twice with 100 µL of ice-cold PBSCMA.

For experiments using only a single yeast transformant (e.g. yeast transformed with pBEVY-pT231 scFv 3.24-GFP), the cells were separated out at a concentration of 10^6^ yeast cells per well. For experiments using two yeast transformants, namely those expressing pBEVY-4420-mCherry in addition to yeast cells expressing pBEVY-pT231 scFv 3.24-GFP, cells were mixed in equal amounts to the desired final concentration. These cells were washed three times with 500 µL of ice-cold PBSCMA and resuspended in PBSCMA at a volume of 50 µL per 10^6^ yeast cells. Yeast cells were then incubated in the wells for 30 minutes at room temperature. After incubation, wells were washed once with 100 µL of ice-cold PBSCMA.

Wells were then washed by dispensing 100 µL of ice-cold PBSCMA to one side of a wall (for example, the east wall side) and aspirating the liquid from the opposite side (the west side). The buffer was then dispensed to a different wall side (for example, the north wall side) and aspirated from the opposite side (the south side). This was repeated for a third time with yet again a different combination of wall sides (for example, northeast to southeast well walls). This whole process was repeated twice again using different combinations of wall sides, making sure to dispense liquid to walls that were aspirated from previously. In other words, if the first round of washing included pipetting from the north to the south side, pipetting from the south to the north side needed to be accounted for. Additionally, we made sure to include wall sides that were not included initially such as the northwest and southwest sides of the well walls. At this point, each well should have been washed once plainly and three times with 3 different combinations of well wall sides. If this protocol is carried out, each well should have been washed (dispensing media and aspirating it) a total of 10 times. After washing, 100 µL of ice-cold PBSCMA were added to the wells and the plate was used for imaging and analysis.

### 96-well Image Acquisition

The automated microscope Keyence BZ-X810 was used for imaging the wells. For each well, a minimum of 4 set points were made and focused within the 96-well plate setting in the BZ-X800 Viewer software. Once the set points were made, the microscope automatically took scanning fluorescence images of the wells. After the images were acquired, they were analyzed by the BZ-X800 Analyzer software. An appropriate threshold was determined manually using an image at the edge of a well and this threshold was applied to all of the images taken during the same day of experiments. All of the images from an individual well were then loaded into the software along with the threshold image, image stitching was turned on, and the software counted the individual cells present. The number reported by the software was used as the “cell count” number in analysis.

### Antibody Labeling of Yeast Displayed scFv

To quantify scFv expression, 2 × 10^6^ yeast cells displaying scFvs were washed twice with PBSA (137 mM NaCl, 2.7 mM KCl, 10 mM Na_2_HPO_4_, 1.8 mM KH_2_PO_4_, and 1 g/L BSA, pH 7.4) and resuspended in 100 µL PBSA with a chicken anti-FLAG tag antibody (Abnova PAB29056, 1:500 dilution). Cells were incubated with the primary antibody for 30 minutes on ice and washed once with 500 µL PBSA. Cells were then stained with either goat anti-chicken IgY Alexa Fluor 647 (ThermoFisher, Cat. No. A21449, 1:200 dilution) or goat anti-chicken IgY Alexa Fluor 488 (ThermoFisher, Cat. No. A11039, 1:200 dilution) in 100 µL PBSA for 30 minutes on ice. Cells were then washed once with 500 µL PBSA and resuspended in 500 µL PBSA before detecting fluorescence.

To evaluate scFv binding to target peptide ligands, 2 × 10^6^ yeast cells were washed twice with PBSA and incubated with the biotinylated phospho-peptide in PBSA at room temperature for 1 hour. The volume of PBSA used for incubation varied depending on the concentration of phospho-peptide that was tested to ensure the peptide to scFv ratio remained constant. After labeling with the peptide, the cells were then stained as described above with streptavidin R-phycoerythrin (ThermoFisher, Cat. No. S866, 1:100 dilution) added with a secondary antibody. Cells were washed and resuspended before detecting fluorescence.

Yeast cell fluorescence was detected using the BD Biosciences LSR Fortessa X-20 flow cytometer (UConn Center for Open Research Resources and Equipment). ‘scFv expression’ and ‘GFP expression’ indicate fluorescence values recorded from the flow cytometer for the FLAG tag staining and GFP fluorescence, respectively. ‘binding/expression’ refers to the fluorescence values from the biotinylated peptide binding divided by the scFv expression.

### Statistical Analysis

Prism 8 (GraphPad) was used for statistical analysis. Statistical tests used for each dataset are described in the figure legends.

## Results

### Development of Yeast Biopanning for a Peptide Antigen

To assess the binding of yeast cells displaying antibody fragments to a peptide antigen target, we developed a biopanning approach (**Fig. 1**) outlined as follows (**Fig. 1a**). First, wells within a 96-well plate were coated with Matrigel to allow for the attachment of HEK293FT cells, which were grown to 100 % confluency. These cells were biotinylated, and then streptavidin was added to the biotinylated HEK293FT cells. Since four individual biotin molecules can bind to one streptavidin molecule, we anticipated there would be leftover streptavidin binding sites to capture biotinylated peptides. After biotinylated peptide was added, yeast cells expressing scFv binders were added.

**Figure 1.**
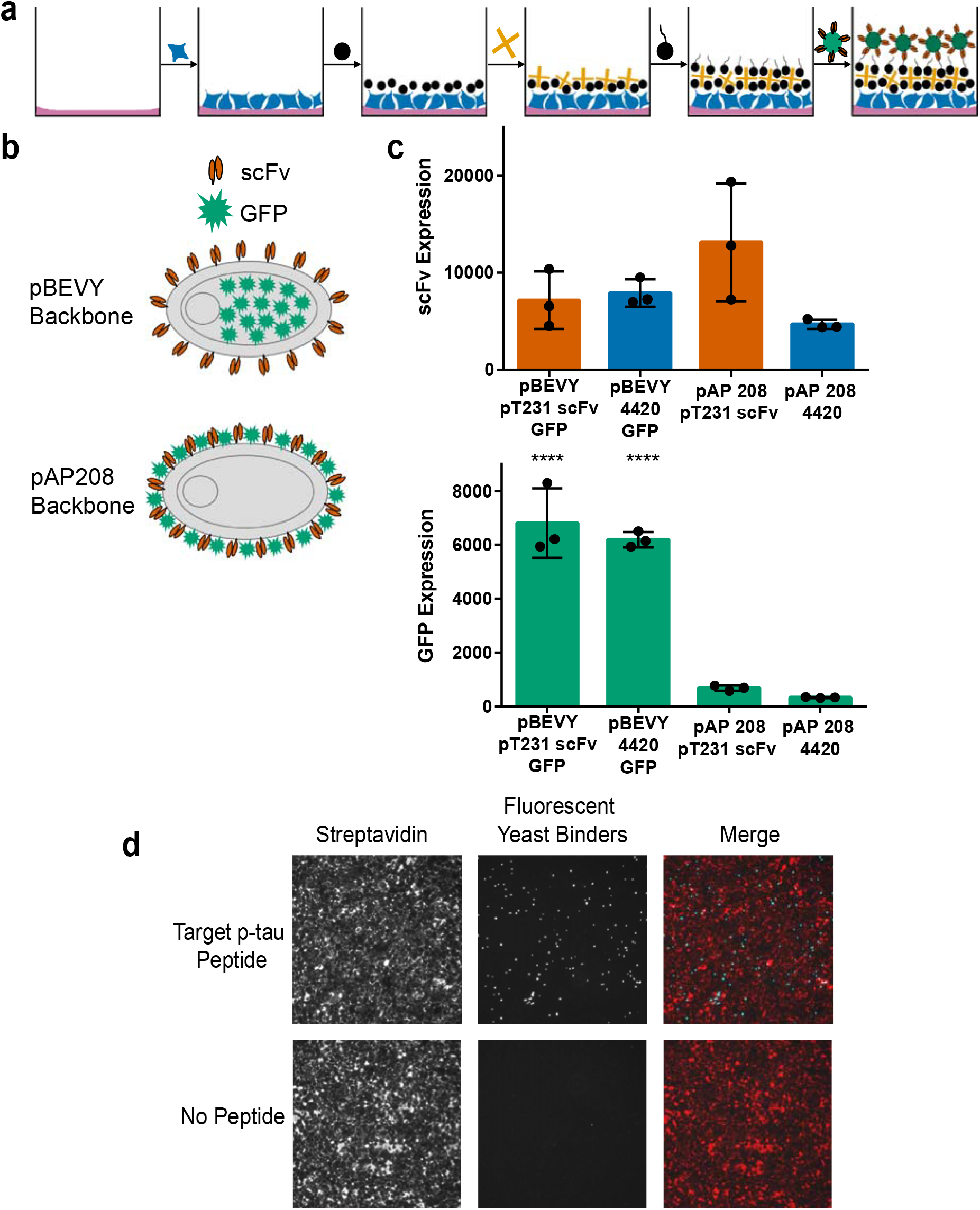
Overview of biopanning method and yeast reagent design. **a**, schematic representation of biopanning method. Wells are first coated with Matrigel and HEK293FT are seeded. The cell surface is then biotinylated, and streptavidin is added. Biotinylated peptides are then added to the wells to be used as target antigens. Yeast cells displaying antibody binders and intracellularly expressing fluorescent protein are added to the wells and used as reporters. **b**, schematic representation of different reporter designs. The pBEVY backbone expresses antibody fragments (scFv) on the yeast cell surface and fluorescent protein (GFP) intracellularly. The pAP208 backbone expresses both scFv and GFP on the yeast cell surface. **c**, comparing expression levels for scFv and GFP. **** *P* ≤ 0.0001 using Tukey’s multiple comparisons test. Otherwise, *P* > 0.05. **d**, representative images indicating presence of streptavidin on HEK293FT cell surface (red) and yeast cells binding to target peptide (cyan).

To facilitate the detection of yeast cells, we designed and tested two different plasmid constructs that allow surface display of scFvs and expression of fluorescent proteins (**Fig. 1b**). Both designs use the Aga2p system (Boder & Wittrup, 1997) for yeast surface display of scFvs. The first design uses bi-directional expression based on the pBEVY backbone (Miller et al., 1998) for intracellular expression of fluorescent proteins (e.g., GFP or mCherry) and surface display of scFvs **(Fig. 1b**). On the other hand, in the second design (pAP208), GFP was fused to the C-terminus of Aga2p (scFv-Aga2p-GFP), enabling its expression on the yeast cell surface (**Fig. 1b**). To these constructs we cloned the pT231 tau scFv, which binds a phosphorylated peptide derived from the human microtubule-associated tau protein (D. Li et al., 2018) or an anti-fluorescein scFv 4420 (Boder & Wittrup, 1997) as a control. We then compared the expression levels of both the scFv and GFP using flow cytometry. We found that there were no significant differences in scFv expression when comparing the two different backbone constructs (**Fig. 1c**). However, when comparing GFP expression levels, there were clear significant differences between the two backbones (**Fig. 1c**). The pBEVY backbone expressed GFP at about 6-7-fold higher levels than the pAP208 backbone (**Fig. 1c**) and was therefore used for the subsequent biopanning experiments.

To quantitatively assess binding, we performed biopanning using yeast cells displaying the high affinity pT231 scFv mutant (pT231 scFv 3.24) in 96-well plates as described in **Fig. 1a**, but with or without the biotinylated phosphorylated tau (p-tau) peptide. After the addition of these yeast binders, the wells were thoroughly washed loosely following the previously described yeast biopanning protocol (Wang et al., 2007) with some modifications (see Methods). Successful biotinylation of HEK293FT cells is shown using streptavidin conjugated with Alexa 647 (**Fig. 1d**). We found clear binding of the yeast cells in wells containing the p-tau peptide, but nearly no binding in wells without the peptide (**Fig. 1d**).

To verify that the yeast cells were selectively binding to the p-tau peptide, we repeated the biopanning experiment with a non-phosphorylated tau peptide in addition to the no peptide control. To comprehensively assess the binding, we used an automated microscope (Keyence BZ-X810) that takes fluorescence images of the entire well and quantified the amount of yeast cells present in each well after washing. These experiments were repeated across three different days with triplicates each day. When analyzing the number of yeast cells in each well, it is clear that they specifically bind to the intended p-tau peptide (**Fig. 2b**). We attribute the relatively small number of cells present in wells with no peptide or non-phosphorylated peptide to the inability to wash out non-binding cells completely.

**Figure 2.**
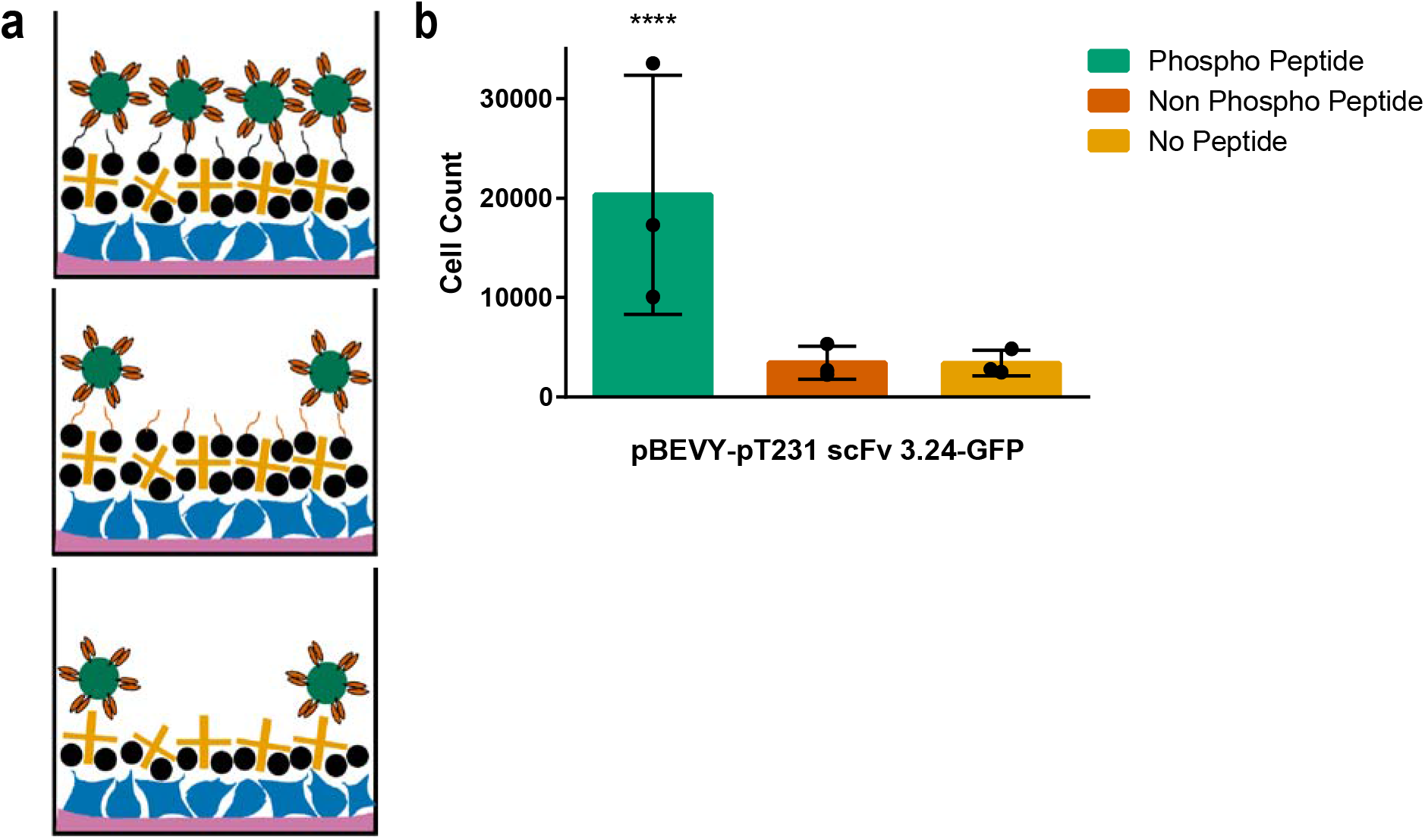
Testing selective binding of the scFv to peptide of interest. **a**, schematic representation showing different control conditions tested. Wells with target peptide (topmost image) are expected to contain more yeast cell reporters than wells with non-target peptide (middle image) and wells without peptide (bottom image). **b**, quantification of yeast cells expressing pT231 scFv 3.24 and GFP present in wells after washing. **** *P* ≤ 0.0001 using two-way ANOVA. Otherwise, *P* > 0.05.

### Biopanning Using an Internal Control Yeast Strain

After observing non-specific signal present in wells with non-phosphorylated peptide and no peptide, we wanted to introduce a more reliable method of capturing non-specific signal. To do this, we designed a second strain of yeast cells that could be used as an internal control for the biopanning experiments (**Fig. 3a**). This control yeast strain expresses mCherry instead of GFP and the anti-fluorescein scFv using the pBEVY plasmid backbone. Using yeast cells expressing this plasmid (pBEVY-4420-mCherry) in conjunction with yeast cells expressing the plasmid described above (pBEVY-pT231 scFv 3.24-GFP) endowed us with the ability to quantify the non-specific signal within each well. The experimental set-up again included wells with p-tau peptide and controls (**Fig. 3b**). Yeast cells expressing pBEVY-4420-mCherry and yeast cells expressing pBEVY-pT231 scFv 3.24-GFP were mixed in equal numbers and added to the wells (**Fig. 3b**). After washing, the wells were imaged and the number of yeast cells in each well was quantified.

**Figure 3.**
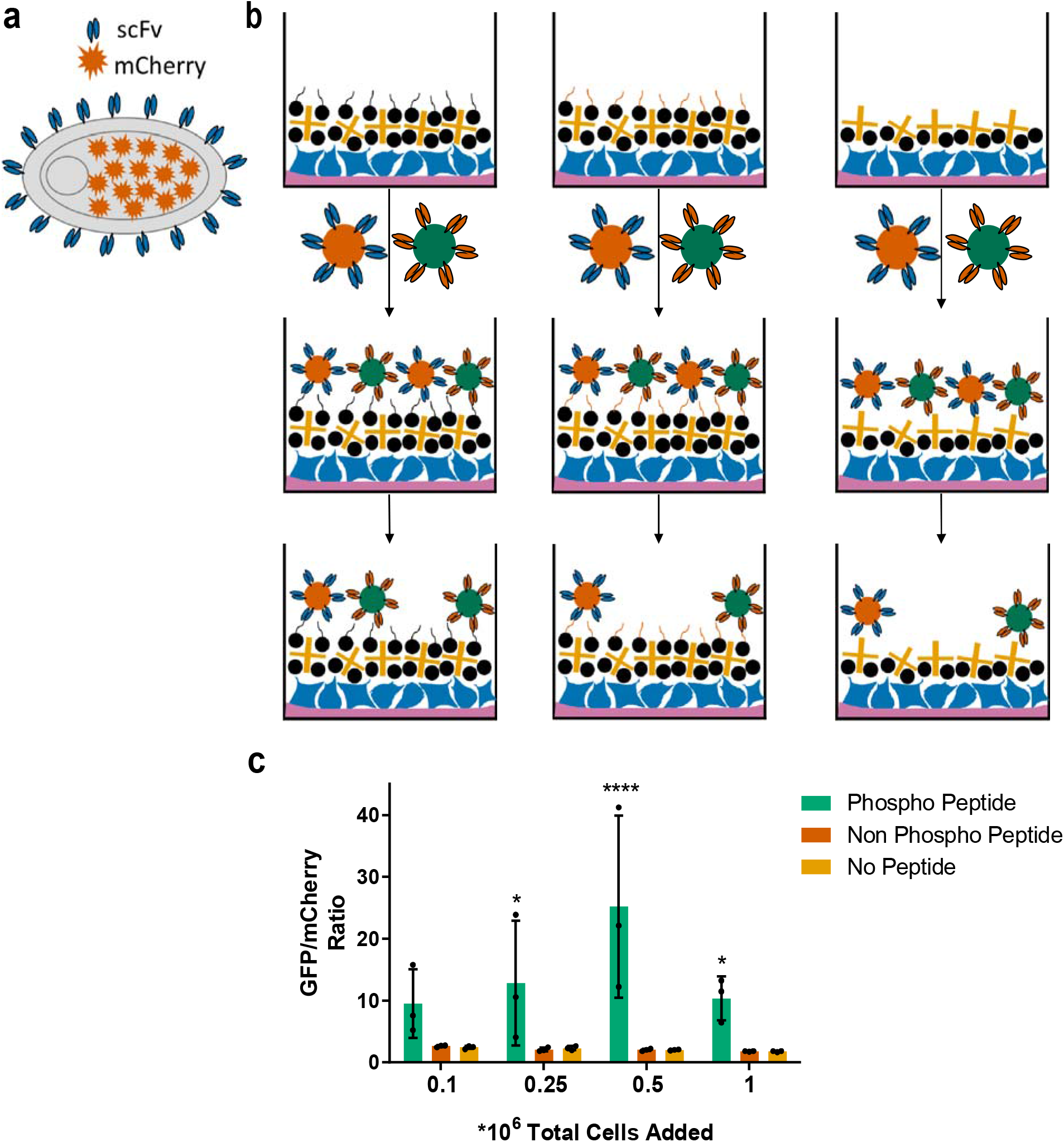
Design and application of internal control yeast cell reporters. **a**, schematic representation of control yeast cell design. Yeast cells display a control scFv on their surface and intracellularly express a fluorescent protein (mCherry). **b**, schematic representation of biopanning method with control yeast cells included. Yeast cells expressing pBEVY-4420-mCherry and pBEVY-pT231 scFv 3.24-GFP are mixed in equal ratio and added to wells containing phospho peptide (left), non-phospho peptide (middle), and no peptide (right). **c**, quantification of the ratio of yeast cells expressing pBEVY-pT231 scFv 3.24-GFP to yeast cells expressing pBEVY-4420-mCherry present in wells after washing. * *P* ≤ 0.05, **** *P* ≤ 0.0001 using Tukey’s multiple comparisons test. Otherwise, *P* > 0.05.

To evaluate if the total number of yeast cells in each well affects the degree of non-specific binding, we tested different cell densities in the biopanning experiment. For each cell density condition tested, the yeast cells expressing the two different plasmid constructs were mixed in equal proportion. After washing, full-well imaging, and cell counting, the ratio of yeast cells expressing pBEVY-pT231 scFv 3.24-GFP to yeast cells expressing pBEVY-4420-mCherry was determined (**Fig. 3c**). These experiments showed that this ratio remained relatively consistent for the control wells, indicating that non-specific binding can be accurately captured by using the internal control yeast strain. Additionally, we observe that adding an initial total amount of 0.5 × 10^6^ yeast yields the best ratio of specific binders to non-specific binders (**Fig. 3c**). Taken together these results show that, while cell density does not affect the degree of non-specific binding, it is important to consider when looking to optimize the best specific to non-specific binder ratio.

### Improved Biopanning Parameters

To further understand the mechanisms of this approach and to achieve the highest possible number of binding yeast cells, we aimed to test and optimize three parameters: the concentration of the biotinylation reagent (NHS-PEG4-Biotin) applied to HEK293FT cells, the concentration of the peptide antigen, and the concentration of the streptavidin reagent that captures biotinylated peptide (**Fig. 4**).

**Figure 4.**
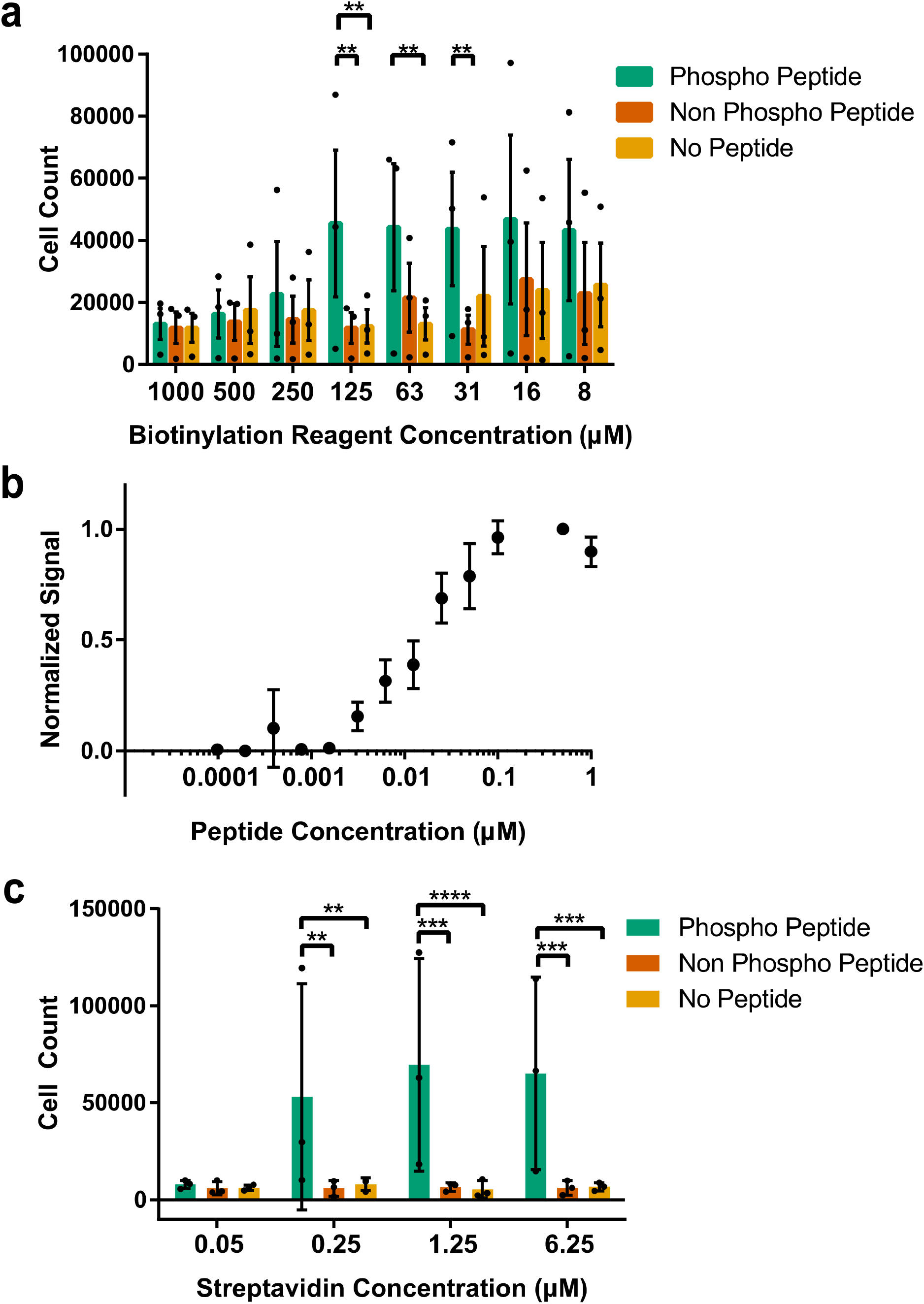
Improving biopanning parameters. **a**, quantification of number of yeast cells expressing pBEVY-pT231 scFv 3.24-GFP present in wells when tested across different biotin concentrations. **b**, normalized “signal” [(# yeast cells, phospho-peptide wells)-(# yeast cells, no peptide wells)]/(# yeast cells, highest concentration of peptide) for a range of peptide concentrations. **c**, number of yeast cells present in wells when tested across different streptavidin concentrations. ** *P* ≤ 0.01, *** *P* ≤ 0.001, **** *P* ≤ 0.0001 using Tukey’s multiple comparisons test. Otherwise, *P* > 0.05.

To analyze the effect of the biotinylation reagent, we tested eight different concentrations ranging from 7.81 µM to 1 mM (**Fig. 4a**). For each of these concentrations, cells expressing pBEVY-pT231 scFv 3.24-GFP were added to wells containing p-tau peptide or controls. After washing and well imaging, the number of cells in each well was determined. Each biotinylation reagent concentration parameter was tested on three separate days with one sample quantified each day. Statistical testing for the differences between target peptide and control peptides for each concentration showed that only three biotin concentrations resulted in significantly higher binding compared to the controls (**Fig. 4a**). At biotinylation reagent concentration of 31.25 µM, there was a significant difference in the cell count of the phosphorylated target peptide when compared with the non-phosphorylated peptide, but not when compared with the cell count of wells without peptide (**Fig. 4a**). At 62.5 µM, there was a significant difference in the cell counts between phosphorylated target peptide and wells containing no peptide, but not when compared to wells containing non-phosphorylated peptide (**Fig. 4a**). Biotinylation reagent concentration of 125 µM proved to be the most promising, showing a significant difference in the cell count of phosphorylated target peptide when compared with both non-phosphorylated peptide and wells without peptide (**Fig. 4a**). For this reason, this reagent concentration was chosen as the standard in subsequent biopanning experiments.

To test the sensitivity of our platform to the peptide target antigen, we tested a range of p-tau peptide concentrations between 0.098 nM to 1 µM (**Fig. 4b**). Each peptide concentration was tested on at least three separate days, with triplicates each day. For analysis of this data, the “signal” (number of yeast cells expressing pBEVY-pT231 scFv 3.24-GFP still present after washing), was normalized. Normalizing the data involved subtracting the yeast cell count present in wells without peptide (taken as background) from the yeast cell count present in wells with target peptide. This number was then normalized to the highest signal from that day of experiments, which was always at least 0.1 µM. From these experiments, we saw no detectable signal from peptide concentrations of 0.098 nM to 1.56 nM (**Fig. 4b**). However, we observed significant binding starting at a peptide concentration of 3.13 nM and apparent signal saturation beginning at a concentration of 0.1 µM (**Fig. 4b**). From these results, a peptide concentration of 0.1 µM was chosen for the purpose of obtaining the strongest signal possible in subsequent experiments.

To further optimize the interactions within this biopanning platform, we tested a range of streptavidin concentrations between 50 nM to 6.25 µM (**Fig. 4c**). For each concentration, cells expressing pBEVY-pT231 scFv 3.24-GFP were added to wells containing p-tau peptide and controls. After washing and full-well imaging, the number of cells in each well was determined and each parameter was tested on three separate days with duplicates each day. Statistical analysis showed that a streptavidin concentration of 50 nM resulted in no significant difference of cells binding to target peptide versus controls. However, each of the three higher streptavidin concentrations (250 nM, 1.25 µM, 6.25 µM) showed a significant difference when comparing cell count in wells with target peptide versus controls (**Fig. 4c**). Furthermore, when comparing the cell count of the p-tau peptide across these three streptavidin concentrations, statistical testing showed no significant differences. For these reasons, a streptavidin concentration of 1.25 µM was chosen to ensure there is enough streptavidin present for the capture of biotinylated peptides.

### Biopanning with a Lower Affinity Variant

Once the most important conditions were optimized, we were interested in testing the performance of biopanning to detect the binding of lower affinity scFvs (**Fig. 5**). Working towards the goal of using this platform to screen pools of antibodies, it is highly important to capture cells displaying antibody fragments with sub-optimal affinities.

**Figure 5.**
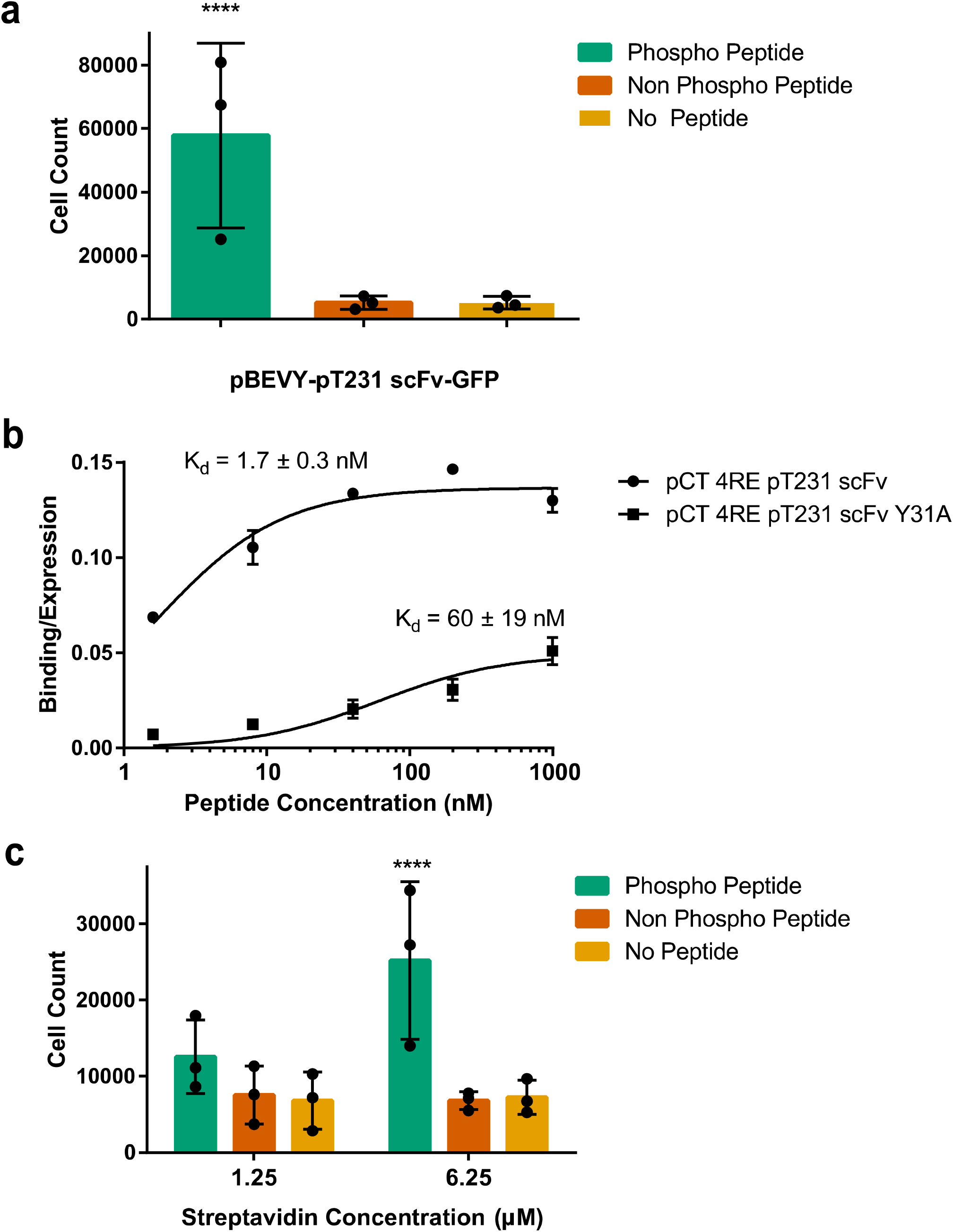
Testing selective binding for lower affinity scFvs to peptide of interest after optimizing parameters. **a**, quantification of number of yeast cells expressing pBEVY-pT231 scFv-GFP present in wells with a biotin concentration of 125 µM, a peptide concentration of 0.1 µM, and a streptavidin concentration of 1.25 µM. **b**, binding/expression curve for wild-type scFv (pCT-4RE-pT231 scFv) and lower affinity mutant (pCT-4RE-pT231 scFv Y25A) measured using flow cytometry. **c**, quantification of yeast cells expressing pBEVY-pT231 scFv Y25A-GFP present in wells after washing. **** *P* ≤ 0.0001 using Tukey’s multiple comparisons test. Otherwise, *P* > 0.05.

Because the pT231 scFv 3.24 used (K_D_ = 200 pM (D. Li et al., 2018)) is a high affinity and specificity mutant of a previously described antibody (pT231 scFv WT, K_D_ = 2.2 nM (Shih et al., 2012)), we first tested this wild-type scFv (**Fig. 5a**). The optimized conditions of biotin (125 µM), peptide (0.1 µM), and streptavidin (1.25 µM) were tested with the wild-type scFv (pBEVY-pT231 scFv-GFP) across three different days with triplicates each day. After washing steps, the wells were imaged and the number of cells in each well were counted and analyzed. These experiments showed a significant difference in the number of cells in p-tau peptide wells when compared to negative controls (**Fig. 5a**).

To test even lower affinity antibodies, we turned our attention to a lower affinity mutant (pT231 scFv containing an alanine point mutation Y31A in the CDR we reported previously (D. Li et al., 2018)) of the original wild type. To compare the affinity of this mutant to the wild type, we performed titrations for yeast cells expressing the wild-type pT231 scFv or the mutant (Y31A) (expressed using the pCT-4RE plasmids cloned previously (D. Li et al., 2018)) with the p-tau peptide at concentrations ranging from 1.6 nM to 1 µM (**Fig. 5b**). These experiments showed that the affinity of pT231 scFv Y31A is approximately 35-fold less than that of pT231 scFv WT, with K_D_ = 60 ± 19 nM (**Fig. 5b**).

The low affinity variant was cloned into the pBEVY backbone co-expressing GFP (pBEVY-pT231 scFv Y31A-GFP) and tested in the biopanning platform (**Fig. 5c**). These experiments were repeated across three different days, with duplicates each day. These experiments showed that a streptavidin concentration of 1.25 µM, which previously appeared to be the concentration at which cell counts leveled off (**Fig. 4c**), no longer resulted in a significant difference between target wells and control wells. From these experiments, we see that further increasing the streptavidin concentration to 6.25 µM does result in significant capture of yeast cells based on phospho-specific antibody binding (**Fig. 5c**). These results indicate that the high streptavidin concentration is necessary when biopanning for low to moderate affinity antibodies, which are often seen in antibody library screening.

## Discussion

We developed a novel method of yeast biopanning against site-specific protein phosphorylations and demonstrated its use against the human tau protein. The approach allows discriminating yeast cells displaying scFvs based on binding to peptides containing a single site-specific phosphorylation. The primary focus of the biopanning approach was on selective yeast capture based on phospho-site specific antibody interaction. We have previously demonstrated the capability to enhance the affinity of phospho-site specific antibodies without sacrificing specificity (D. Li et al., 2018), but clonal selection strategies based on phospho-specificity is still critically lacking. Here we demonstrated that the biopanning is highly selective to yeast cells expressing phospho-specific scFvs, and only to peptides that contain the phosphorylated residue. The bi-directional expression plasmids to display scFvs and intracellularly express fluorescent protein reporters allowed efficient counting of interacting yeast cells, and in-well monitoring of non-specific binding. This allowed us to improve biopanning conditions, such as streptavidin and biotinylation reagent concentration. Finally, we show that scFvs with moderate affinities (K_D_ = 60 nM) can be detected, well within the range of antibodies identified from scFv libraries using yeast display biopanning (Wang et al., 2007; Zorniak et al., 2017).

Notably, the streptavidin concentration needed to be increased by 5-fold to capture the yeast cells displaying the lower affinity scFv mutant (pT231 scFv Y31A) (**Fig. 5c**). In addition to having lower affinity, the Y31A mutant also showed lower binding signal normalized to expression level (**Fig. 5b**), indicating that a greater fraction of displayed scFv may not be functional compared to the wild-type scFv. This will reduce the surface density of available scFvs that interact with the target, which may be why greater streptavidin concentration was required. Such sub-optimal scFv folding is a likely scenario in heterologous antibody fragment libraries, and therefore further demonstrates the robustness of the yeast biopanning method for detecting phospho-site specific interaction.

The monolayer of HEK293FT cells on which the biotinylated peptides are immobilized seem superfluous but our initial attempts to biopan without them were unsuccessful (data not shown). The mammalian cell surface provides a matrix in which proteins are embedded that can be biotinylated. This complex structure may provide higher avidity interactions that allow yeast cell binding. For library screening, a negative selection step will be necessary to eliminate antibodies that bind to non-targets, including the mammalian cell surface antigens, biotin, and streptavidin. Negative selections have been already demonstrated using yeast biopanning to identify cell-type selective binders (Zorniak et al., 2017).

Although we have demonstrated this capability using a single phosphorylation site in tau, we anticipate this method will be widely applicable to phospho-specific antibody characterization and screening. Due to the fact that protein phosphorylation sites tend to be located in disordered regions (Iakoucheva et al., 2004; Nicolaou et al., 2021), synthetic peptides containing phosphorylated residues have been extremely effective antigens for generating phospho-site specific antibodies suitable for a wide range of applications, including various cell and tissue labeling, immunoblotting, and immunoassays. With this new capability to biopan against phospho peptides, identification of monoclonal antibodies will be greatly aided by rapid characterization of phospho-site specific binding of clones after immunization. Moreover, screening various existing yeast surface display libraries, including the human scFv (Feldhaus et al., 2003), camelid nanobodies (McMahon et al., 2018), and other binders (Hackel et al., 2008; Kruziki et al., 2015) will be possible. Combined with subtractive panning methods to eliminate non-specific binders, this approach has great potential for identifying phospho-site specific clones. In particular, we anticipate to identify antibodies against tau phosphorylation sites that need specific antibodies (Arbaciauskaite et al., 2021).

## Acknowledgements

This work was funded by grants NSF 1706743, NIH 1R21NS111358-01A1, and Alzheimer’s Association 2019-AARG-NFT-640971. M.A. was also supported by the GE innovation fellowship.

